# Genetically distinct parallel projection populations from ventral hippocampus to prefrontal cortex

**DOI:** 10.1101/2023.02.18.529052

**Authors:** Candela Sánchez-Bellot, Andrew F. MacAskill

## Abstract

The ventral hippocampus is proposed to perform its multitude of roles via segregated populations of neurons identified by either genetic makeup, downstream projection, or their combination. Recently we described two parallel projections from ventral hippocampus to prefrontal cortex, segregated across the radial axis of the CA1 and subicular areas. These two populations had distinct afferent and efferent connectivity and distinct influence of approach avoidance behaviour. In this study, we extend these data by performing RNA sequencing of each population of neurons. We find that these two populations have multiple genes that are differentially expressed. These genes correspond both to genes classically thought to be distributed across the radial axis such as *Calbindin 1* and *Pcp4*, but also to more unexpected genes including postsynaptic scaffolds and GABA receptor subunits. Notably, a number of genes differentially expressed across the two populations were associated with the development of mental illness, suggesting an imbalance in the function of these two pathways in disease may be an interesting area for future research. Together, these data reinforce the dissociation of function of projections to prefrontal cortex across the radial axis of the ventral hippocampus, and provide multiple targets for both the genetic and functional dissociation of these roles.

## INTRODUCTION

The ventral hippocampus, and in particular its output structure the CA1 and subiculum is thought to be composed of multiple, parallel populations of neurons that can be separated based on their genetic identity or downstream projection target^1^. This parallel structure is proposed to allow the hippocampal circuit to perform multiple distinct calculations, ranging from goal directed behaviour, to spatial navigation and memory^2,3^.

One of the most heavily investigated projections is that from ventral CA1 and subiculum to prefrontal cortex (vH-PFC)^4^. This projection has been shown to be crucial for a wide array of different behaviours, including goal directed spatial navigation, working memory, approach avoidance behaviour and threat extinction^4–8^. Consistent with this wide-ranging role, while many projections form hippocampus (for example to nucleus accumbens and amygdala) overlap strongly with a specific genetically defined population of neurons^9^, those projecting to PFC are genetically diverse^10^, suggesting that this projection may be composed of more than one subpopulation of neurons^9^. Consistent with this, we recently described two parallel populations of neurons in vH that both project to PFC but are distributed across the radial axis of ventral CA1 and subiculum^4^. The first, arising from the superficial layers preferentially targets interneurons in PFC and promotes exploratory behaviour. In contrast the second, located in the deep layers, contacts fast spiking interneurons and pyramidal neurons, and promotes avoidance behaviour. We found that the widely used superficial genetic marker *Calbindin 1* could be used to separate these two populations, suggesting that they may represent distinct genetically defined populations – consistent with previously reported variability in vH-PFC genetic identity^9^. However, to what extent these two populations have differential gene expression remains unknown.

In this study we performed RNA sequencing of fluorescently identified PFC projecting vH neurons from both the superficial and deep layers. We found numerous differentially expressed genes, ranging from scaffold and adhesion proteins, to GABA receptor subunits and trafficking proteins. By comparing our sequencing data to previous freely available datasets, we show that superficial and deep vH-PFC neurons correspond to distinct, previously identified genetically defined populations of vH neurons. Together, these data emphasise the distinct nature of the vH projection to PFC, and show that the deep and superficial layers are not only distinct at a cellular, circuit and behavioural level^4^, but are also distinct at the molecular level.

## RESULTS

### The projection form vH to PFC is composed of two parallel populations distributed across the radial axis

We first replicated our previous finding showing that projections to PFC from vH are segregated across the radial (superficial to deep) axis^4^. In previous work we have characterised these two projection populations using either cholera toxin, AAVretro or retrobead labelling^4^. Therefore, we took this opportunity to investigate the distribution of vH-PFC neurons labelled instead with rabies virus – to ensure that the two-layered distribution was not due to some otherwise unidentified tropism of labelling. To do this we utilised a non-pseudotyped, G-deleted rabies virus expressing a red fluorescent mCherry cassette. This virus can infect presynaptic terminals and thus infect neurons that project to the area of injection. However, the lack of the rabies glycoprotein in the viral package ensures the virus can only trace back one synaptic connection – precluding polysynaptic spread of the virus. We injected this virus into the PFC of 3 mice (**Figure 1a**), and after 9 days prepared coronal sections of both PFC and vH. Consistent with our previous work^4^, we found a marked segregation of labelled neurons in vH into two populations – one along the superficial layer and one in the deep layer (**Figure 1b**).

**Figure 1.**
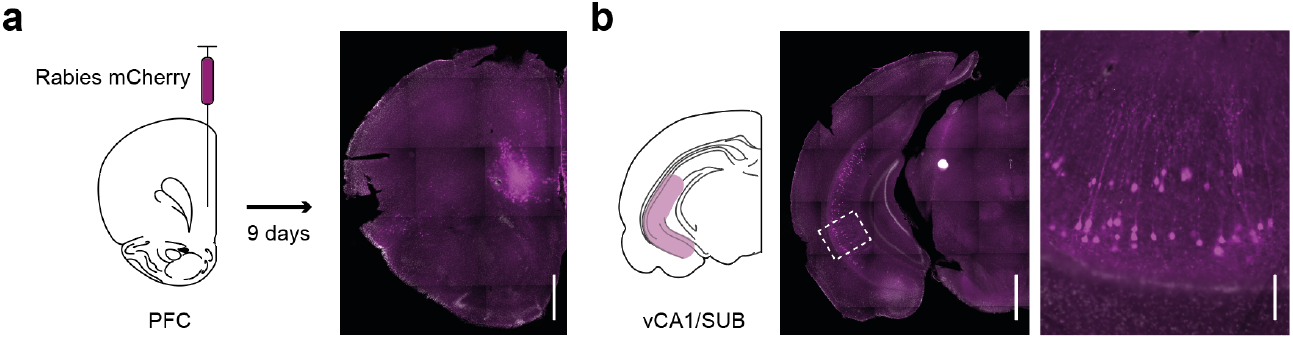
Retrograde rabies labelling reveals two populations of vH neurons that project to PFC distributed across the radial axis. Injection of G-deleted rabies expressing mCherry into the PFC labels PFC neurons (**a**), but also neurons in vH that project to PFC (**b**). Zoom in (**b**) shows vCA1 / subiculum border with two clear populations of neurons, one superficial layers, one in deep layers. Image representative of data from 3 mice. Scale bars 1 mm, 1 mm, 200 μm.

### Differential RNA expression in each population of PFC projecting vH neurons

We next investigated whether these two populations of neurons may have differential gene expression. To do this we injected 4 mice bilaterally in PFC with retrobeads (**Supplementary Figure 1**) – bright, fluorescent latex microspheres that are taken up by presynaptic terminals and transported retrogradely to label afferent neuronal somas. 2 weeks later, we confirmed prominent segregation of fluorescence across the radial axis in acute slices of vH (**Figure 2b**). Guided by this fluorescence, we then dissected the two layers of labelled vH neurons (e.g. **Figure 2c**), and isolated single cells from these sections using a combination of enzymatic and manual dissociation (see methods). Cell suspensions were then sorted using fluorescence assisted cell sorting – using red fluorescence to identify and select retrobead labelled neurons, and DAPI labelling to exclude unviable, permeabilised cells – before being pooled and processed for population RNA sequencing (**Figure 2d**, see methods). This approach obtained high-read-depth, high-quality transcriptomes, and data have been deposited in the National Center for Biotechnology Information (NCBI) Gene Expression Omnibus under GEO: GSE225512.

**Figure 2.**
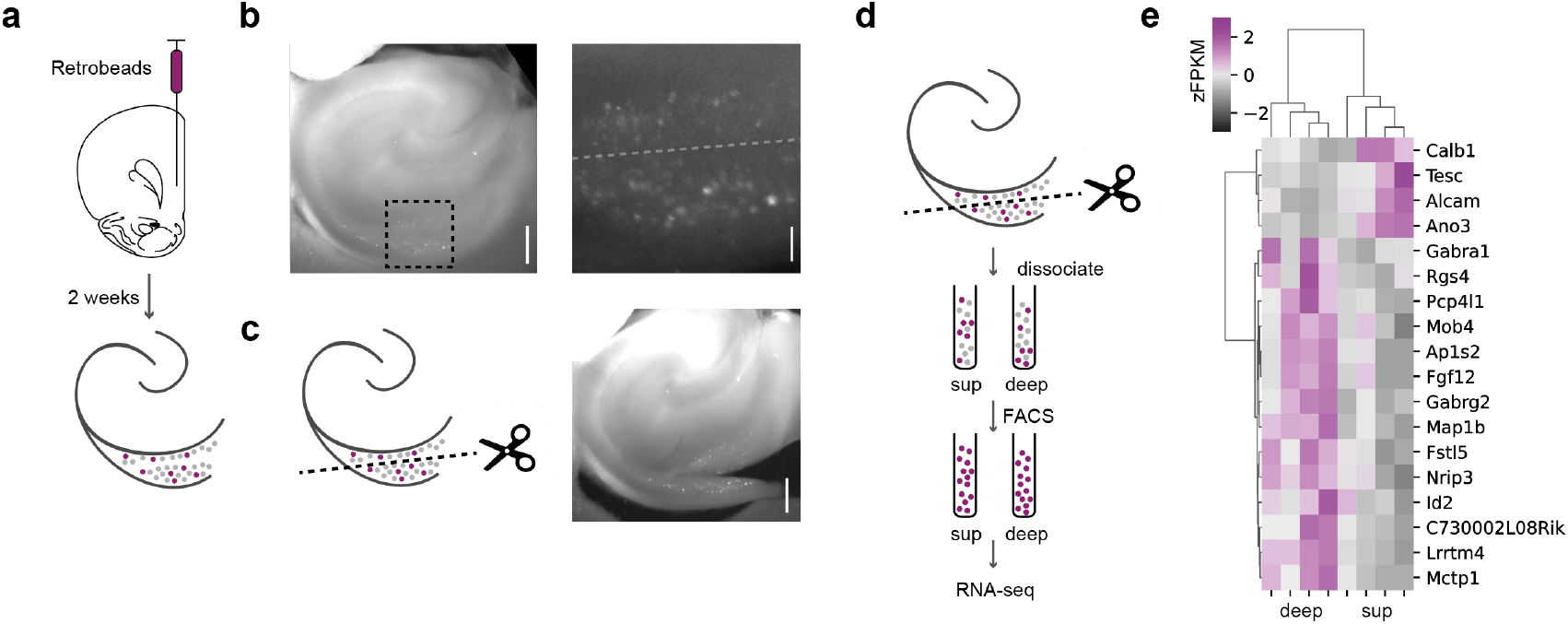
Dissection and RNA sequencing of superficial and deep vH-PFC projection populations. **a**. Injection of retrobeads into the PFC labels PFC-projecting neurons in vH. **b** Example acute slice showing dual-layer distribution of fluorescently labelled neurons. **c**. The same acute slice after a cut was made to separate the two layers for separate analysis. **d**. Analysis workflow for dissociation, FACS purification and RNA sequencing (see methods for details). **e**. Heatmap showing differentially expressed genes across two layers as zFKPM, ordered using Bray-Curtis similarity. Only genes which passed multiple comparison testing are shown for clarity. Scale bars 1 mm.

We first confirmed that our sorting approach had successfully isolated excitatory neuronal populations that would be expected from retrograde labelling^4^. We compared expression of excitatory neuronal genes such as CaMKii, Synapsin, and vGlut1 - which showed robust expression - with markers for oligodendrocytes, astrocytes, microglia and interneurons – which all showed very low levels of expression (**Supplementary Figure 2**). Together this suggests that our sorting approach successfully isolated excitatory pyramidal neurons.

We next investigated how different genes varied across each population of vH neurons that project to PFC. Comparing reads from superficial and deep groups across mice, we found 18 genes that were robustly differentially expressed across the two layers (**Figure 2e**), 4 enriched in the superficial population, and 14 enriched in the deep population (This increases to 31 genes if slightly less stringent statistical criteria are applied – **Supplementary Figure 3**). These genes included classic markers of superficial vH neurons that have been widely used such as *Calb1*^4,11^, and well-known deep layer markers such as *PCP4*^9,12^. In addition, we also found a number of other genes such as specific GABA-A receptor isoforms (*GABRA1* and *GABRG2*), and synaptic scaffold proteins (*LRRTM4*) that were enriched in the deep population. Consistent with proposed distinct connectivity within and across the radial axis in vH^4,13–15^

We next wanted to validate potential differentially expressed genes. To do this we investigated how expression of candidate genes differed across the radial axis in the Allen Brain Atlas dataset^16^. We found that the majority of genes identified in our screen displayed marked segregation across the radial axis, and were enriched in either the superficial or deep layers as predicted by our sequencing results (**Figure 3a-d**), suggesting that expression of these genes could dissociate cells across the radial axis.

**Figure 3.**
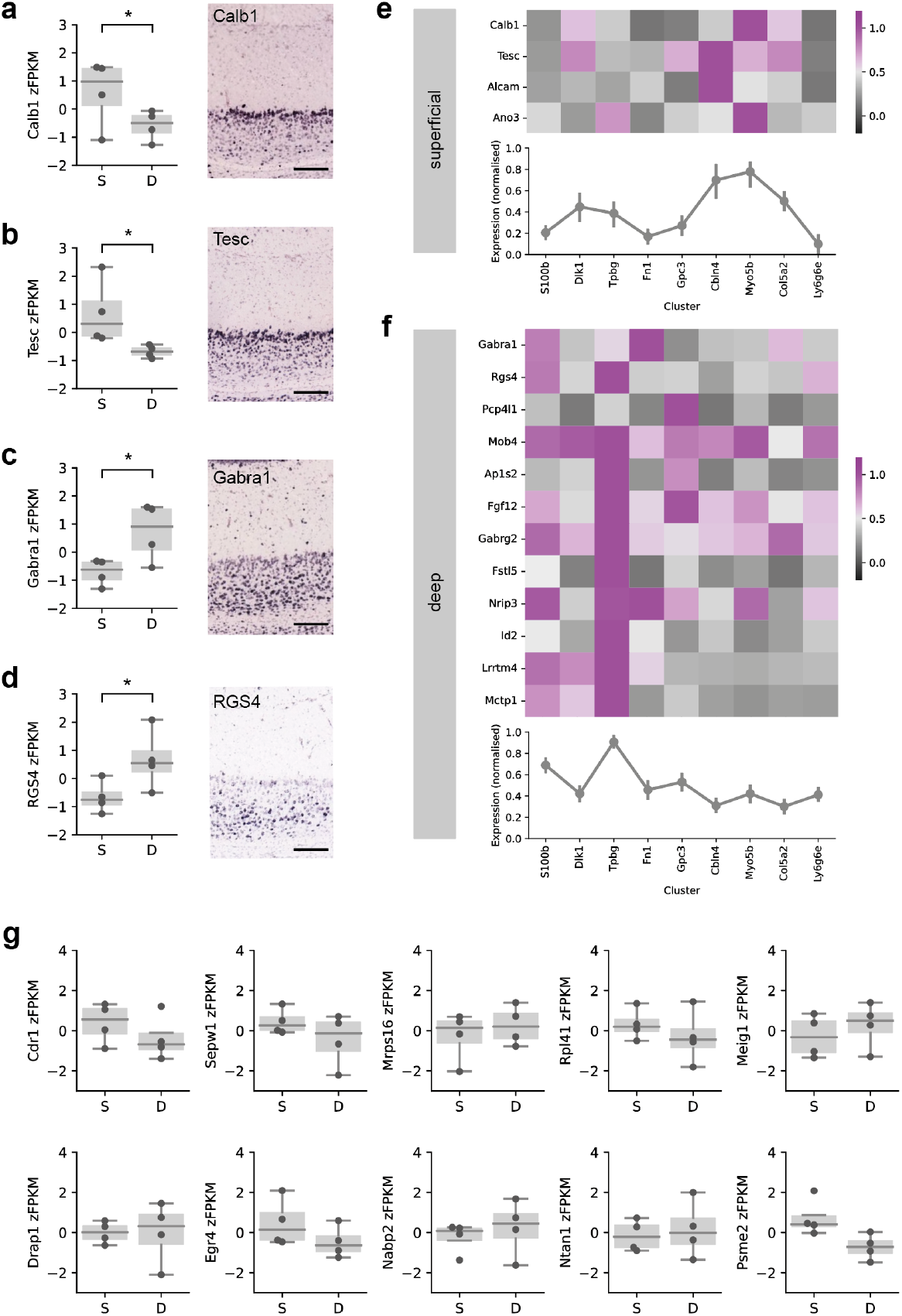
Validation of hits and comparison to other datasets. **a-d**. Validation of 4 example genes. *Left* shows zFKPM from sequencing datasets, *right* shows coronal section of vH subfields. In all cases both show *Calb1* and *Tesc* are enriched in superficial layers, while *Gabara1* and *RGS4* are enriched in deep layers. Scale bars 1 mm, 1 mm, 200 μm. **e-f**. Comparison to data from ref.9 showing that superficial enriched transcripts are prevalent in *Cbln4, Myo5b* and *Col5a2* clusters, while deep transcripts are prevalent in Tpbg clusters. **g**. Relative expression from sequencing of 10 genes shown to have enriched expression specifically in vH-PFC neurons in ref.10. Note that none of these genes show differential expression across the radial axis.

Finally, we wanted to compare our results to recent descriptions of vH-PFC RNA expression. First, recent data has defined vH neurons as clusters of genetically distinct neurons in CA1 and subiculum^9,10^. Notably, in contrast to projections to nucleus accumbens and amygdala that seem to corresponded to only one or two genetic clusters, PFC projecting neurons were found to be remarkably distinct, and be composed of a number of different genetic clusters, differentiated by the expression of marker genes *S100b, Dlk1, Tpbg, Fn1, Gpc3, Cbln4, Myo5b, Col5a2* and *Ly6g6e*. Therefore, we reasoned that the distinct populations of neurons projecting to PFC across the radial axis may account for some of this previously unexplained variability in expression. To investigate this, we compared the expression of each of our candidate genes to each previously identified cluster^9^. We found that genes expressed by superficial located PFC projecting vH neurons were enriched in clusters associated with *Clbn4, Col5a2* and *Myo5b* (**Figure 3e**). In contrast, genes expressed in deep located PFC projecting neurons were more generally expressed across all clusters, but specifically enriched in the cluster associated with *Tpbg* (**Figure 3f**). Therefore vH-PFC neurons, when separated across the superficial and deep layers can be assigned into distinct previously identified populations of neurons.

Second, a recent study found that vH-PFC neurons were markedly distinct from other vH projection neurons, mainly through differential expression of 24 genes, of which 11 - *Cdr1, Sepw1, Mrps16, Rpl41, Meig1, Mrpl52, Drap1, Egr4, Nabp2, Ntan1, Psme2* - were upregulated specifically in vH-PFC neurons^10^. We therefore investigated to what extent these transcripts were differentially expressed across the superficial and deep layers. While each of these transcripts were present in detectable amounts in our experiment, interestingly none of these genes were differentially expressed across the two layers (**Figure 3g**). This suggests that different sets of genes differentiate vH-PFC neurons from other vH neurons, than differentiate vH-PFC neurons across the radial axis.

## DISCUSSION

In this study, we have shown that neurons in vH that project to PFC from either the deep or superficial layers of the radial axis have unique molecular signatures. This is consistent with previous work showing marked variability across the radial axis on vH^9,10,16–20^ and also marked variability within neurons that project to PFC^9^. Moreover, it is consistent with recent findings showing deep and superficially located neurons in hippocampal CA1 and subiculum have distinct properties^21–23^, and that particularly in the case of the projection to PFC their roles in circuit function and control of behaviour are remarkably distinct^4^.

The genetic variability of vH neurons has been highlighted by two recent studies^9,10^. Both these studies found that across different projection populations in vH (projecting to e.g. PFC, NAc, amygdala and hypothalamus), there was marked and consistent genetic specialisation. Moreover, it was recently shown that this specialisation was much more extreme in neurons projecting to PFC^10^ – suggesting that these form a unique subclass of vH neurons that are far more distinct from other projections. Notably, in our study we recapitulated these sequencing results and found the majority of mRNA transcripts previously associated specifically with PFC projecting neurons, including *Cdr1, Rpl41, Meig1*, and *Ntran1*. However, it is interesting to note that each of these transcripts were not different across the radial axis – with equal expression in both superficial and deep layers. This suggests an interesting dichotomy – where PFC neurons express a set of genes that render them genetically distinct en masse, but where each layer expresses a separate subset of genes that differentiates them across the radial axis. Further work looking at how other projection populations differ across the radial axis would allow an appreciation of how much these differences in expression across the radial axis are specific to PFC projection populations, or are a more general feature of radial axis specialisation irrespective of projection population.

Interestingly, when sequenced at a single cell level, when compared to other projection populations, vH-PFC projection neurons are also much more internally variable^9^. This variability can be clustered into 9 distinct populations, that were proposed to represent genetically distinct subpopulations in vH – identified by the genes *S100b, Dlk1, Tpbg, Fn1, Gpc3, Cbln4, Myo5b, Col5a2* and *Ly6g6e*. Therefore, we investigated the distribution of genes differentially expressed over the radial axis in PFC projecting neurons across these distinct clusters. We found that transcripts enriched in superficial vH-PFC neurons were highly represented in three of these clusters – *Col5a2, Cbln4* and *Myo5b*. Interestingly, *Col5a2* and *Cbln4* clusters have been associated with superficial layers in intermediate and ventral vH respectively, while *Myo5b* is associated with neurons in the proximal CA1/subiculum border – consistent with the dorsoventral and proximal localisation of superficial vH-PFC neurons^4,24^. In contrast, deep layer vH-PFC neurons were more variably distributed, but enriched in the cluster associated with *Tpbg*, which again has previously been associated with deep layer neurons^9^. Together this analysis confirms the distinct genetic identity of superficial and deep neurons, and suggests that by separating vH-PFC neurons into superficial and deep layer populations, much of the previously defined variability in their genetic makeup can be reduced.

A key feature of this analysis was that when compared to single cell datasets, the deep layer had expression of genes across multiple clusters, albeit at different levels. Interestingly, our recent work also showed that deep layer neurons seemed to be composed of a number of different neurons, and had an increased probability that neurons would project to more than one downstream area^4^. Therefore, an interesting possibility is that the superficial layer represents a single genetically defined population, while the deep layer may be further subdivided. Future work to understand what additional factors could be utilised to probe this potential variability will help clarify if this is the case.

Finally, it is notable that a high proportion of genes differentially expressed across the superficial and deep layers of vH-PFC neurons are risk factors for mental health disorders. For example, RGS4 and LRRTM4 and Alcam have been proposed as risk factors for bipolar disorder and schizophrenia^25–28^, but almost all genes identified in our screen have been associated with mental health disorders, and in particular in mood disorders and schizophrenia. We recently found that superficial and deep layers are poised to exert tight control over PFC excitatory : inhibitory balance^4^. Therefore, the differential impact of these genes across the two layers of vH-PFC neurons is potentially consistent with a proposed key role for excitation : inhibition balance in mental health^29^. Future work investigating how the two layers of vH-PFC neurons are affected in preclinical models of mental health disorders can elucidate this.

Overall, we have shown that vH neurons that project to PFC are composed of two layers spread across the radial axis, and that these two populations are genetically distinct. Coupled with our previous work showing dramatic differences in circuit and behavioural roles of these populations^4^, this work provides a genetic basis for understanding their function in more detail.

## METHODS

### Surgery

Stereotaxic injections were performed on 6-8 week old mice anaesthetised with isofluorane (4 % induction, 1 – 2 % maintenance) as injections carried out as previously described^4^. Briefly, the skull was exposed with a single incision, and small holes drilled in the skull directly above the injection site. Injections are carried out using long-shaft borosilicate glass pipettes with a tip diameter of ∼ 10 - 50 μm. Pipettes were back-filled with mineral oil and front-filled with ∼ 0.8 μL of the substance to be injected. A total volume of 140 - 160 nL of virus or retrobeads was injected at each location in ∼ 14 or 28 nL increments every 30 s. The pipette was left in place for an additional 5 min to minimize diffusion and then slowly removed. Injection coordinates for infralimbic PFC were as follows (mm relative to bregma): ML: ± 0.4, RC: + 2.3, and DV: - 2.4. After injection, the wound was sutured and sealed, and mice recovered for ∼30 min on a heat pad before they were returned to their home cage. Animals received carprofen in their drinking water (0.05 mg / ml) for 48 hrs post-surgery as well as subcutaneously following surgery (0.5 mg / kg).

### Histology

Mice injected with rabies virus were perfused with 4% PFA (wt/vol) in PBS, pH 7.4, 9 days after surgery, and the brains dissected and postfixed overnight at 4ºC as previously described^4^. 70 μm thick slices were cut using a vibratome (Campden Instruments) in the coronal plane. Slices were mounted on Superfrost glass slides with ProLong Gold antifade mounting medium (Molecular Probes). NucBlue was included to label gross anatomy. Imaging was carried out with a Zeiss Axio Scan Z1, using standard filter sets for excitation/emission at 365-445/50 nm and 545/25-605/70 nm. Raw images were analysed with FIJI.

### RNA Sequencing

Four 6 - 8 week old mice were injected with red retrobeads in PFC (see *surgery*). After 2 weeks, acute, 400μm thick transverse hippocampal slices were prepared. Mice were anaesthetized with a lethal dose of ketamine and xylazine, and perfused intracardially with ice-cold external solution containing (in mM): 210 sucrose, 7 glucose, 25 NaHCO_3_, 1.25 NaH_2_PO_4_, 2.5 KCl, 1 Na^+^ ascorbate, 3 Na^+^ pyruvate, 7 MgCl_2_ and 0.5 CaCl_2_, bubbled with 95% O_2_ and 5% CO_2_. Slices were cut in this solution and then transferred to artificial cerebrospinal fluid (aCSF) containing (in mM): 119 NaCl, 25 glucose, 25 NaHCO_3_, 1.25 NaH_2_PO_4_, 3 KCl, 1 Na^+^ ascorbate, 4 Na^+^ pyruvate, 1 MgCl_2_ and1.3 CaCl_2_, bubbled with 95% O_2_ and 5% CO_2_, and incubated for 30 minutes at 35°C. aCSF contained TTX (100 nM) to block sodium channels, NBQX (20 μM) and APV (50 μM) to block AMPA and NMDA receptor transmission respectively, and Pronase (1 mg / ml) to disrupt the extracellular matrix. This incubation was followed by a further 30 mins wash in aCSF without Pronase. Following incubation, slices were placed in a sylgard-filled chamber under a dissection stereomicroscope (Leica MZ16FA) equipped with fluorescence and transmitted light. The ventral CA1 / subiculum was separated from the rest of the slice, and a single cut was made to separate superficial and deep cell layers, guided by the fluorescence of PFC-projecting cells. Superficial and deep excised sections from each animal were then pooled (4 samples of superficial and deep sections per animal), and tissue was manually triturated for dissociation using three glass Pasteur pipettes of decreasing tip diameter. The resulting suspension was mixed with DAPI (Molecular Probes) to monitor cell health, and then separated using FACS (UCL FACS facility) to isolate red fluorescent (PFC-projecting), and viable (DAPI negative) cells in superficial and deep layers. Cells were sorted directly into buffer for SMART-Seq v4 Ultra Low Input RNA Kit (Clontech Laboratories, Inc.). Subsequent processing was carried out by the UCL Genomics sequencing facility. Briefly, SMART (Switching Mechanism at 5’ End of RNA Template) technology produces full-length PCR amplified cDNA starting from low input (as low as a single cell). 12 cycles of PCR were used to generate cDNA from lysed cells. The amplified cDNA was checked for integrity and quantity on the Agilent Bioanalyser using the High Sensitivity DNA kit. 200pg of cDNA was then converted to sequencing library using the Nextera XT DNA protocol (Illumina, San Diego, US). This uses a transposon able to fragment and tag double-stranded cDNA (tagmentation), followed by a limited PCR reaction (12 cycles) adding sample specific indexes to allow for multiplex sequencing. For sequencing, libraries to be multiplexed in the same run are pooled in equimolar quantities, calculated from Qubit and Tapestation fragment analysis. Samples were sequenced on the NextSeq 500 instrument (Illumina, San Diego, US) using a 43bp paired end run, generating approximately 200M read pairs in total. Fragments Per Kilobase Million (FPKM) scores were calculated from the raw data using TopHat and Cufflinks via Illumina Basespace software. Only raw FKPM scores over 10 were included. Due to the manual nature of the dissection, for identification of robust markers for the two layers, only hits where the standard deviation across samples in the two conditions was 1 or less were included in final analysis. Statistical significance was compared across superficial and deep samples using false discovery rate correction and hits were defined as those where q < 0.05. The distribution along the radial axis of the hippocampus of a selection of hits were verified using in situ hybridisation data from the Allen Brain Atlas^16^.

## ACKNOWLEDGEMENTS

We thank Marco Beato for providing rabies virus. We thank members of the MacAskill laboratory for helpful discussion. A.F.M.was supported by a Sir Henry Dale Fellowship jointly funded by the Wellcome Trust and the Royal Society (grant number 109360/Z/15/Z) and by a UCL Excellence Fellowship. C.S.B. was supported by the Wellcome Trust 4-year PhD in Neuroscience at UCL (grant number 206074/Z/17/Z).

## DATA AVAILABILITY

Raw and processed RNA-seq datasets have been deposited in the National Center for Biotechnology Information (NCBI) Gene Expression Omnibus under GEO: GSE225512.

## SUPPLEMENTARY FIGURES

**Supplementary Figure 1.**
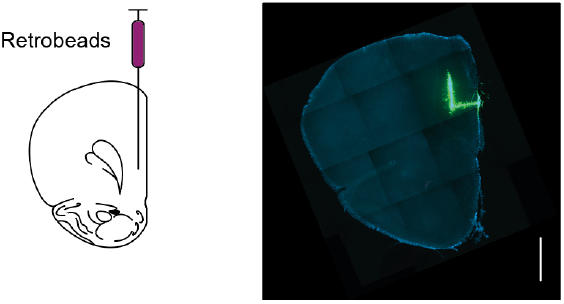
Example retrobead injection into PFC.

**Supplementary Figure 2.**
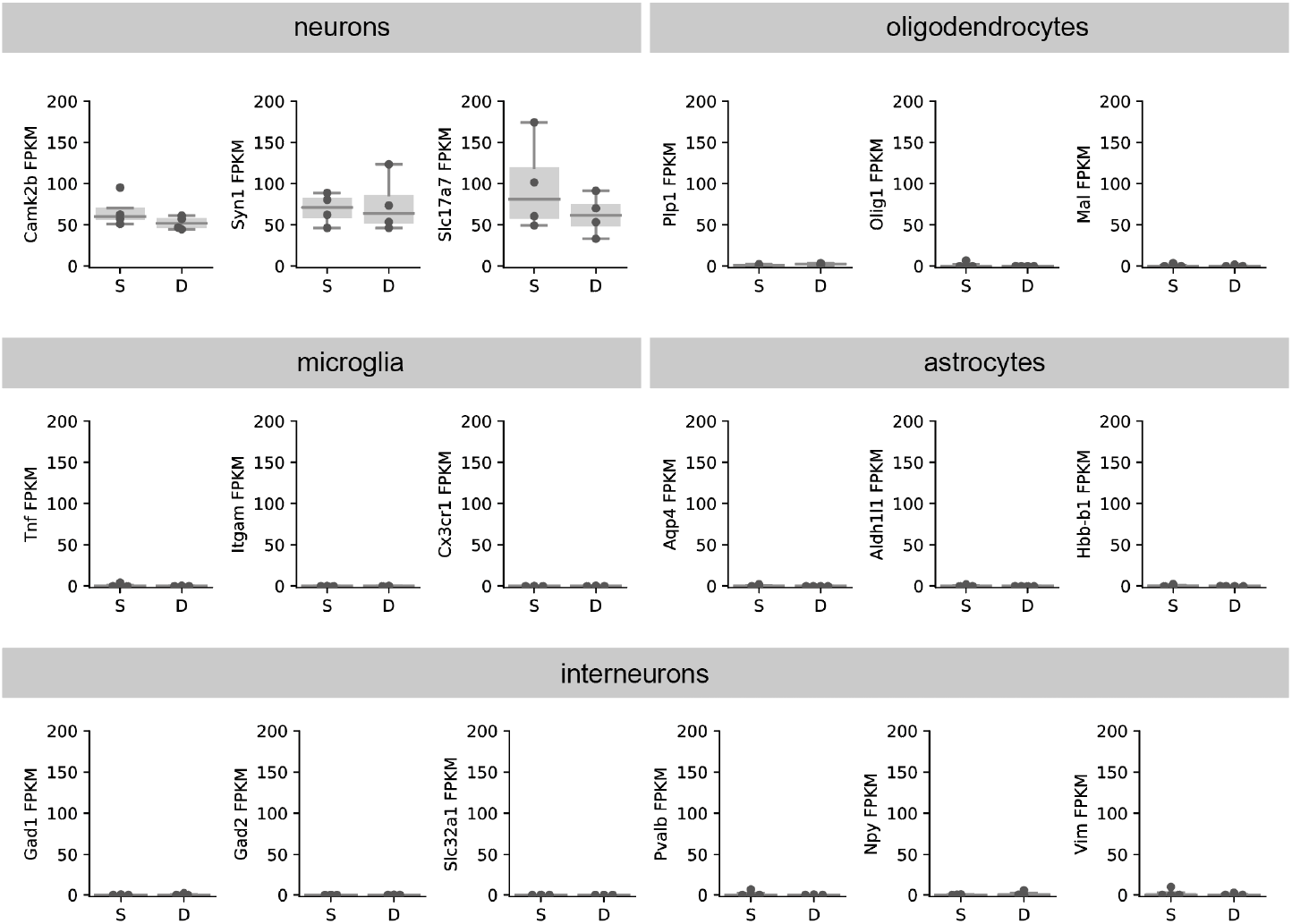
Sequencing of vH-PFC neurons in both layers is enriched in excitatory neuronal transcripts. Raw FKPM values for excitatory neuron, oligodendrocyte, microglia, astrocyte and inhibitory interneuron transcripts, showing that vH-PFC sequencing is enriched for excitatory transcripts.

**Supplementary Figure 3.**
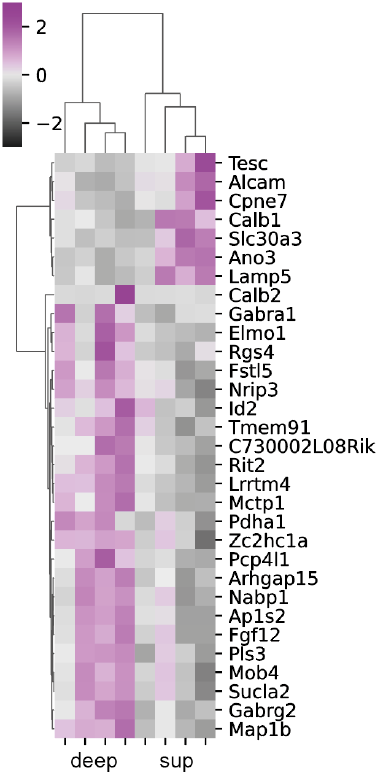
Differentially expressed genes identified with a less stringent q < 0.1 criterion. Graph as in Figure 2e.

## Notes

### Competing Interest Statement

The authors have declared no competing interest.

